# Dual RNA-seq reveals no plastic transcriptional response of the coccidian parasite *Eimeria falciformis* to host immune defenses

**DOI:** 10.1101/117069

**Authors:** Totta Ehret, Simone Spork, Christoph Dieterich, Richard Lucius, Emanuel Heitlinger

## Abstract

**Background:** Parasites can either respond to differences in immune defenses that exist between individual hosts plastically or, alternatively, follow a genetically canalized (“hard wired”) program of infection. Assuming that large-scale functional plasticity would be discernible in the parasite transcriptome we have performed a dual RNA-seq study of the full lifecycle of *Eimeria falciformis* using infected mice with different immune status (e.g. naïve versus immune animals) as models for coccidian infections.

**Results:** We compared parasite and host transcriptomes (dual transcriptome) between naïve and challenge infected mice, as well as between immune competent and immune deficient ones. Mice with different immune competence show transcriptional differences as well as differences in parasite reproduction (oocyst shedding). Broad gene categories represented by differently abundant host genes indicate enrichments for immune reaction and tissue repair functions. More specifically, TGF-beta, EGF, TNF and IL-1 and IL-6 are examples of functional annotations represented differently depending on host immune status. Much in contrast, parasite transcriptomes were neither different between Coccidia isolated from immune competent and immune deficient mice, nor between those harvested from naïve and challenge infected mice. Instead, parasite transcriptomes have distinct profiles early and late in infection, characterized largely by biosynthesis or motility associated functional gene groups, respectively. Extracellular sporozoite and oocyst stages showed distinct transcriptional profiles and sporozoite transcriptomes were found enriched for species specific genes and likely pathogenicity factors.

**Conclusion:** We propose that the niche and host-specific parasite *E. falciformis* uses a genetically canalized program of infection. This program is likely fixed in an evolutionary process rather than employing phenotypic plasticity to interact with its host. In turn this might (negatively) influence the ability of the parasite to use different host species and (positively or negatively) influence its evolutionary potential for adaptation to different hosts or niches.

## BACKGROUND

The term plasticity describes the ability of genetically identical organisms to display variable phenotypes, e.g., via different developmental or metabolic programs. So called reaction norms govern how a particular genotype is translated into a phenotype depending on environmental stimuli [1]. The presence of predators is known to alter, e.g., developmental programs of genetically identical prey animals to produce different phenotypes (reviewed in [2]). Infections by pathogens are known to alter host phenotypes: in fact all non-constitutive immune reactions can be regarded as a manifestation of plasticity [3]. Hence, to understand the outcomes of parasitic infections and host-parasite interactions the concept of plasticity is useful.

The reciprocal effect of the within-host environment on parasite phenotypes, i.e. plasticity, is less studied, especially in parasites of animals. For many parasite species it remains unclear whether differences in pathology are due to parasites’ genotypic or phenotypic (plastic) differences, the latter resulting from host-parasite interactions, e.g., host immune responses.

An exception are Nematode infections (reviewed by [4]), in which for example worm length [5] and other aspects of morphology [6], or developmental timing [7] has been shown to vary with host genotype. However, it is unclear to which extent such differences are passively imposed on the parasite or whether they are responses with functional relevance as an adaptation of the parasite expressing observed phenotypes.

Only recently have transcriptomes been used to investigate plasticity in “infection programs”, which parasites induce as a response to host signals. Since gene expression is orchestrated by the genetic makeup of an organism, plasticity in transcription – when it occurs – is likely to be an adaptation which allows the parasite to react on host stimuli and to produce an altered phenotype. We here distinguish between such plastic (responsive) transcription programs and what is sometimes referred to phenotypic plasticity, which then is a “passive” phenotypic change imposed on the parasite without being controlled at the transcriptional level. A perceivable example could be reduced growth due to “mechanical” impact, e.g., limited space. In a Nematode, the presence of phenotypic plasticity has for example been shown to lack a transcriptional basis [8], and can therefore be regarded “passive”. In contrast, unicellular *Entamoeba* spp. infections of variable pathogenicity (i.e. phenotypic plasticity) manifested also in transcriptional differences under various in vitro conditions [9]. Among apicomplexan parasites, different infection programs with distinct transcriptional profiles have been proposed: in *Plasmodium* spp., the parasite’s transcriptome is distinct in different mouse genotypes (BALB/c and C57BL/6) and tissues within one genotype [10], hence demonstrating the capability for plasticity in this parasite. Similarly and even more closely related to *Eimeria* spp., the coccidian *Toxoplasma gondii* forms dormant tissue cysts (bradyzoites), a process induced by and depending on the host environment [11], and involving large changes in parasite transcriptomes [12]. In addition, *T. gondii* is capable of infecting all studied warm-blooded vertebrates and all nucleated cells in those animals [13] suggesting parasite plasticity in different host environments also in the tachyzoite stage.

*E. falciformis* is an intracellular parasite in the phylum Apicomplexa, which comprises more than 4000 described species [14]. Prominent pathogens of humans are found in this phylum, such as *T. gondii,* the causative agent of toxoplasmosis, *Plasmodium* spp., causing malaria, and *Cryptosporidium* spp., which cause cryptosporidosis. Coccidiosis is a disease of livestock and wildlife caused by coccidian parasites which are dominated by > 1,800 species of *Eimeria* [14]. The genus is best known for several species which are problematic for the poultry industry [15]. *E. falciformis* naturally infects wild and laboratory *Mus musculus,* and its genome is sequenced and annotated making it a useful model for studying *Eimeria* spp. [16]. The parasite has its niche in the cecum and upper part of colon, mainly in the cells of the crypts [17,18]. This monoxenous parasite goes through asexual (schizogony) and sexual reproduction, which results in the host releasing high numbers of oocysts approximately between day six and 14 after infection. When a mouse ingests *E. falciformis* oocysts, one sporulated oocyst releases eight infective sporozoites inside the host, which infect epithelial crypt cells. Within the epithelium, merozoite stages form in several rounds of asexual reproduction, followed by gamete formation and sexual reproduction, within the same host. Schizogony takes place approximately until day six and then gametes form and sexual reproduction takes place, resulting in unsporulated oocyst shedding. Schizogony is not completely synchronous; the exact number of schizogony cycles is unclear and could vary naturally [17,19]. There is evidence for a genetic predisposition of *Eimeria* spp. to perform different numbers of schizogony cycles, as parasites can be selected to become “precocious”, completing the lifecycle faster with a reduced number of schizogony cycles [20,21]. Such results have not been obtained for *E. falciformis,* and similarly, it is not known whether such parasite programs are plastic and can also be triggered by exogenous stimuli, such as host immune responses.

*Eimeria* spp. generally induce host protection against reinfection [19, 22–24] and T-cells seem to play a major role [25,26]. In responses to *E. falciformis* infection of laboratory mice, IFNγ is upregulated [18]. In an IFNγ-deficient mouse host model which displays larger weight losses and intestinal pathology but also lower oocyst output, the wild-type phenotype was recovered by blocking IL-17A and IL-22 signaling [27]. These studies demonstrate that adaptive immunity clearly plays a role in limiting the reproductive success of *Eimeria* spp. infection, but effects on the parasite, apart from reproductive output, remain poorly understood. It is an open question whether the parasite is passively impacted or responds, e.g., via changes in its transcriptome, to changes in the host immune response.

We used a “dual RNA-seq” approach, i.e., we simultaneously assessed the transcriptomes of host and parasite in biological samples containing both species [28–32]. Applying this to an infection of *E. falciformis* in the mouse, we produced host and parasite transcriptomes from the same samples, tissue, and time-points. We describe and analyze host and parasite mRNA profiles at several time-points post infection and contrast transcriptomes of naïve and challenge infected wild-type mice to hosts with strong deficiency in adaptive immune responses. This approach allows us to screen transcriptional changes which may be involved in host-parasite interactions for plasticity to alterations in the host immune system. We hypothesize that changes in the parasite transcriptome would be indicative of a plastic response allowing for functionally altered infection programs.

## RESULTS & DISCUSSION

### Immune competent hosts induce protective immunity against *E. falciformis* infection

To investigate *E. falciformis* development throughout the lifecycle in a natural mouse host (NMRI mice) dual transcriptomes were produced at 3, 5, and 7 days post infection (dpi). We also investigated parasite development and transcriptomes in a mouse strain which is severely limited in adaptive immune responses (*Rag1*^−/−^; “immunocompromised” hereafter) with *Rag1*^−/−^ and the respective isogenic background strain (C57BL/6 as control) at day 5 post infection. To further elucidate host immune responses and parasite sensitivity to host immunity, we also challenge infected all mouse groups (i.e. infected after recovery of a first infection; see Methods) and sampled at the same time-points as in naïve mice.

Infections showed drastically decreased oocyst output (Figure 1A and B) in immune competent hosts undergoing a second, challenge infection compared to naïve animals infected for the first time (Mann–Whitney test, in NMRI, n = 12, U = 32, p = 0.004; in C57BL/6, n = 24, U= 111, p = 0.008). Similarly, a strong reduction of parasite 18S rRNA in the challenge infection down to 3.5% of the amount measured in naïve hosts was detected in reverse transcription quantitative PCR (RT-qPCR) in NMRI hosts (Figure 1C). The model inferring this had a good fit (R^2^ = 0.94) and the change of the intercept for challenged compared to naïve hosts was highly significant (t = −6.71; p < 0.001). Differences in the slope were not significant (t = −1.522; p = 0.15), indicating that the amount of parasite material on 3 days post infection is sufficient to explain a linear increase until 7 days post infection. Overall this data is in line with the strong reduction of oocyst shedding seen in challenge infected immune competent mice, and suggests that the host immune defense disturbs the parasite during gamogony or oocyst formation. Further, these results do not give support to drastic changes in the parasite’s “infection program” and rather suggests a non-plastic lifecycle progression.

**Figure 1.**
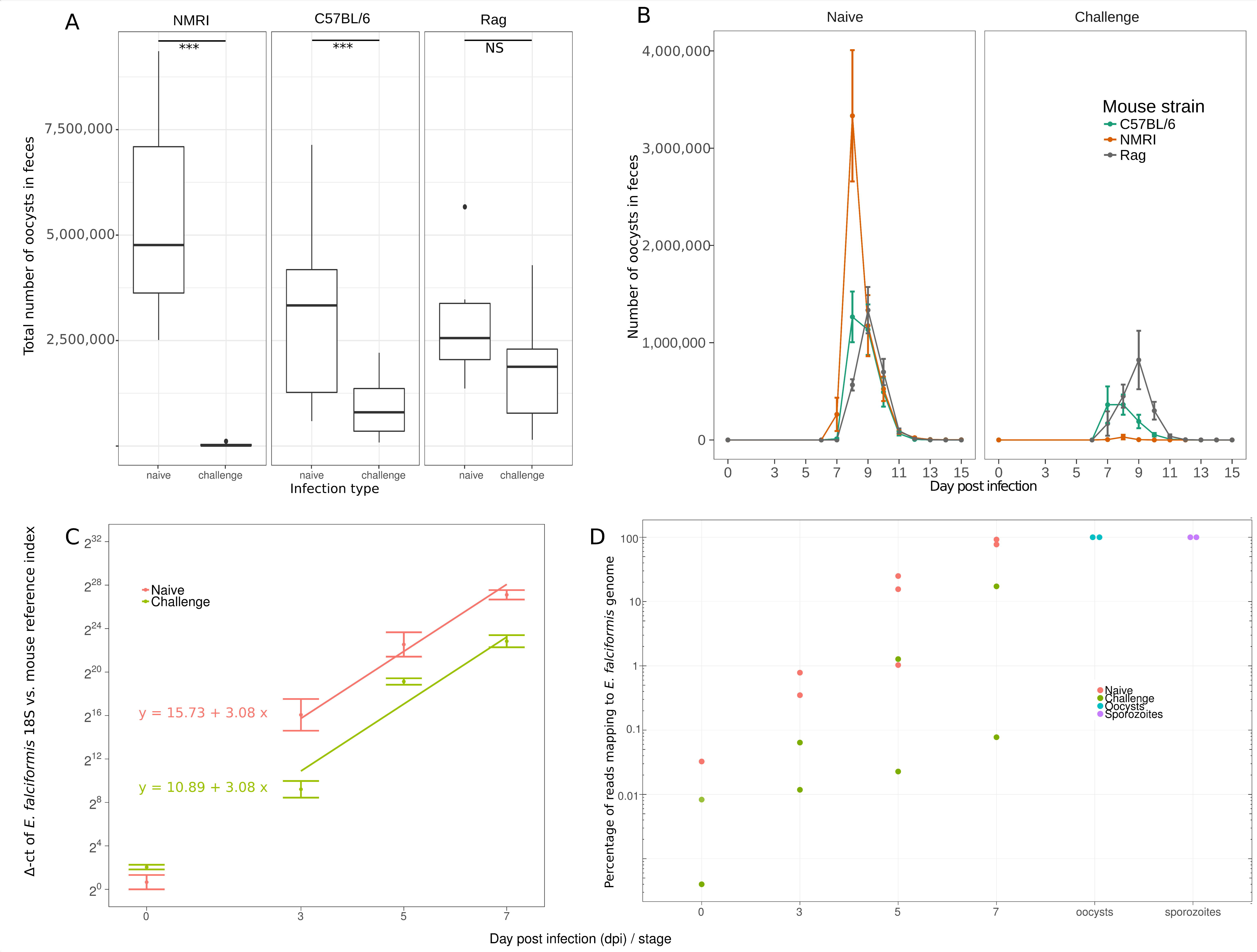
Oocyst output and changes in intensity of *E. falciformis* infection in mouse. Oocyst counts in naïve and challenge infection are shown for three different mouse strains. For infection of naïve NMRI 150 oocysts were used, for challenge infection 1500 oocysts. For C57BL/6 and *Rag1*^−/−^ mice 10 oocysts were used in each infection. A) Overall output of shed oocysts and B) shedding kinetics are depicted. C) RT-qPCR data of *E. falciformis* 18S in NMRI mice displays an increase in parasite mRNA over the course of infection. Significantly less parasite 18S transcripts (normalized against host transcripts of house-keeping genes) were detected in challenge infected mice. Formulas and prediction lines are given for linear models. D) The percentage of parasite mRNA detected by RNA-seq increases during infection (shown for NMRI). More mRNA is detected in naïve mice compared to challenge infected mice. Sporozoites and oocysts contained ~100% parasite material.

In contrast, in immune deficient mice no significant difference in parasite reproductive success (Figure 1A) was observed between naïve and challenge infection (Mann–Whitney test; n = 24, U = 96, p = 0.10). Both in the immunocompromised and immune competent animals, however, all mice had cleared the infection by day 14. We thereby note that *E. falciformis* infection is self-limiting also in mice without mature T- and B-cells, however with a delayed peak of oocyst shedding in immune deficient hosts (Figure 1B).

### Parasite and host dual transcriptomes can be assessed in parallel

We found the increase in parasite numbers over time after infection to also be reflected by the proportion of *E. falciformis* mRNAs sequenced in the combined pool of transcripts from host and parasite (for NRMI mice in Figure 1D). Using mRNA from infected cecum epithelium we demonstrate that even early in infection (3 dpi, during early asexual reproduction) there is sufficient parasite material to detect parasite mRNAs in the pool including host mRNAs, and to quantify individual host and parasite mRNA abundance (Table 1). The number of total (host + parasite) read mappings for individual replicates ranged from 25,362,739 (sample Rag_1stInf_0dpi _rep1) to 230,773,955 (NMRI_2ndInf_5dpi_rep1).

**Table 1.**
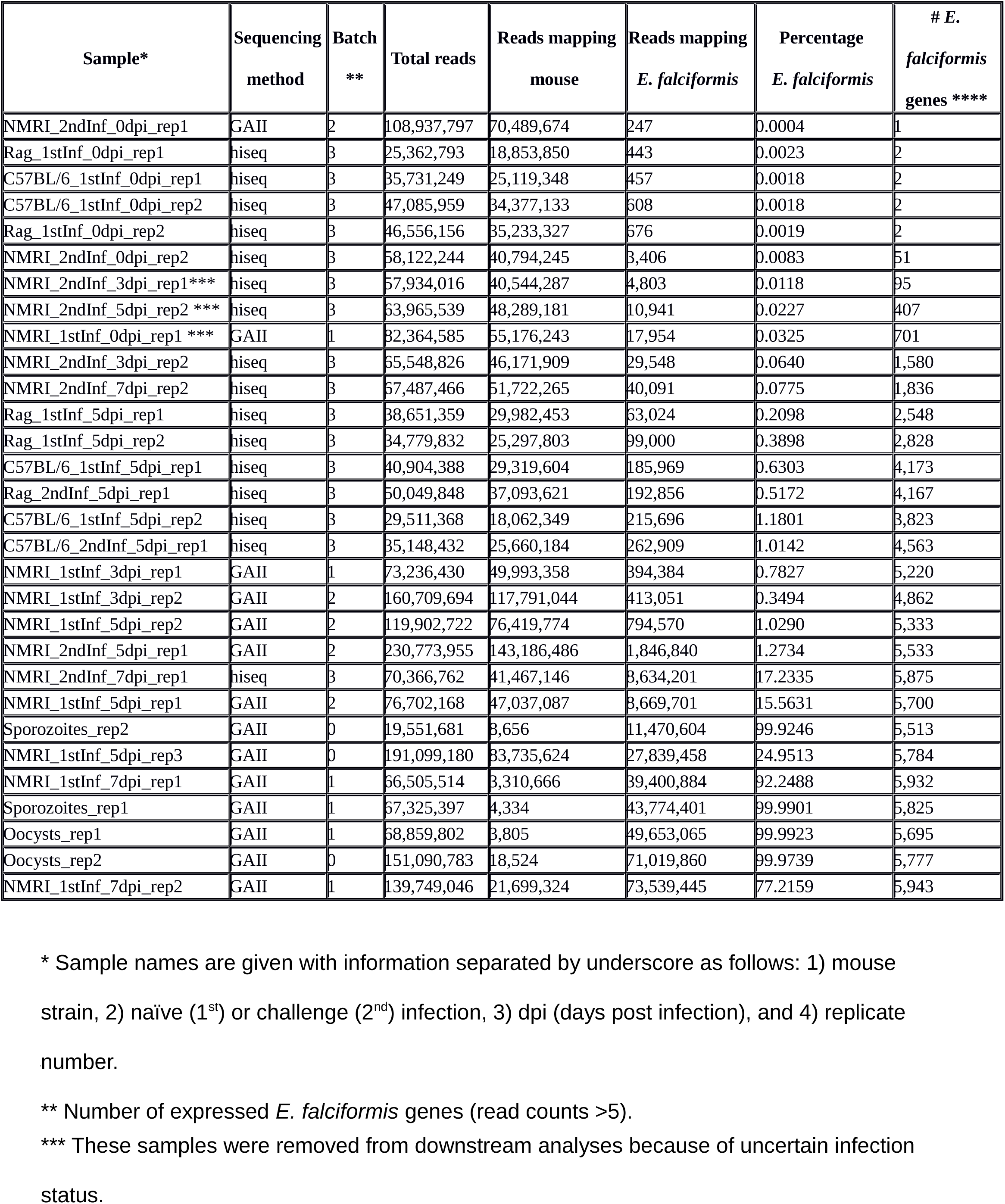
Summary of data per sample, sorted according to number of reads mapping to the *E. falciformis* genome.

We did not detect bias in overall mRNA abundance patterns induced by, e.g., sequencing technologies (batch effects) using a multivariate technique (multidimensional scaling). Efficient normalization was confirmed in that samples with large differences in parasite read proportions show similar transcriptome signatures (Figure S1). This normalization also resulted in unimodal distributions of read numbers (Figure S2) in agreement with negative binomial distributions assumed for statistical modeling and testing.

Remarkably, on day 7 post infection, the day before oocyst shedding peaks, samples from infected naïve mouse epithelium contained 77% and 92% parasite mRNA, i.e., drastically more mRNA from the parasite than from the host (Figure 1D and Table 1). Our transcriptomes for these late infection samples are in agreement with previously published microarray data from mice infected with *E. falciformis* [18], as log2 fold-changes at our 7 days post infection versus controls correlated strongly – for given mRNAs – with log2 fold changes at 6 days post infection versus controls in that study (Spearman’s σ = 0.72, n = 9017, p < 0.001; Figure S3). Considering both biological differences in the experiments, such as exact time-points for sampling, and technical differences between the two methods, this correlation confirms the adequacy of using dual RNA-seq for assessing the host transcriptome in the presence of large proportions of parasite mRNA. Below, we first describe changes in the mouse transcriptome and suggest possible mechanisms at play. Variance in host transcriptome changes upon infection constitutes a potential environmental stimulus for parasites to react on, as addressed later.

### The mouse transcriptome undergoes large changes upon *E. falciformis* infection

We here show that upon infection with *E. falciformis,* which induces weight loss (Figure S4) and intestinal pathology in mice, the host transcriptome undergoes drastic changes affecting more than 3000 individual mRNA profiles significantly (edgeR; glm likelihood-ratio tests corrected for multiple testing, false discovery rate [FDR] < 0.01, see below). Statistical testing for differential abundance between infected and uninfected mice revealed that differences in mRNA abundance were more pronounced (both in magnitude and number of genes affected) at the two later time-points post infection (Table 2 and Figure 2A). 325 mRNAs were differently abundant (FDR < 0.01) between controls and 3 dpi, 1,804 mRNAs between controls and 5 dpi, and 2,711 mRNAs between controls and 7 dpi. This leads to a combined set of 3,453 transcripts responding to infection. Differentially abundant mRNAs early in infection (3 and 5 dpi) were not a mere subset of genes differentially abundant later in infection (7 dpi; Figure 2A), which would be the case if the same genes were regulated throughout infection. Instead, the transcriptional profile of the mouse changes more fundamentally with different genes varying in abundance late compared to early in infection.

**Table 2.** Number of mouse and *E. falciformis* mRNAs significantly differentially abundant in different comparisons (Contrasts). Empty cells indicate that comparison is not applicable.

| Contrast | Number of *E. falciformis* mRNAs<br>with FDR < 0.01 | Number of mouse mRNAs<br>with FDR < 0.01 |
| --- | --- | --- |
| NMRI 7 dpi vs. uninfected control |  | 2,711 |
| NMRI 5 dpi vs. uninfected control |  | 1,804 |
| NMRI 3 dpi vs. NMRI 7 dpi | 1,399 | 1,322 |
| C57BL/6 5 dpi vs. uninfected control |  | 919 |
| NMRI 7 dpi naïve vs NMRI 7 dpi challenge | 0 | 857 |
| NMRI 5 dpi vs. NMRI 7 dpi | 2,084 | 732 |
| *Rag1*<sup>-/-</sup> vs C57BL/6 |  | 362 |
| NMRI 3 dpi vs ctrl |  | 325 |
| C57BL/6 5 dpi naïve vs C57BL/6 5 dpi challenge | 0 | 175 |
| *Rag1*<sup>-/-</sup> 5 dpi vs control |  | 42 |
| NMRI 3 dpi naïve vs NMRI 3 challenge | 1 | 18 |
| NMRI 3 dpi vs. NMRI 5 dpi | 103 | 0 |
| NMRI 5 dpi vs. oocysts | 3,691 |  |
| Sporozoites vs. oocysts | 3,532 |  |
| NMRI 3 dpi vs. oocysts | 3,303 |  |
| NMRI 7 dpi vs. oocysts | 3,202 |  |
| NMRI 7 dpi vs. sporozoites | 2,663 |  |
| NMRI 5 dpi vs. sporozoites | 1,726 |  |
| NMRI 3 dpi vs. sporozoites | 1,705 |  |
| NMRI control vs. C57BL/6 control | 13 |  |

**Figure 2.**
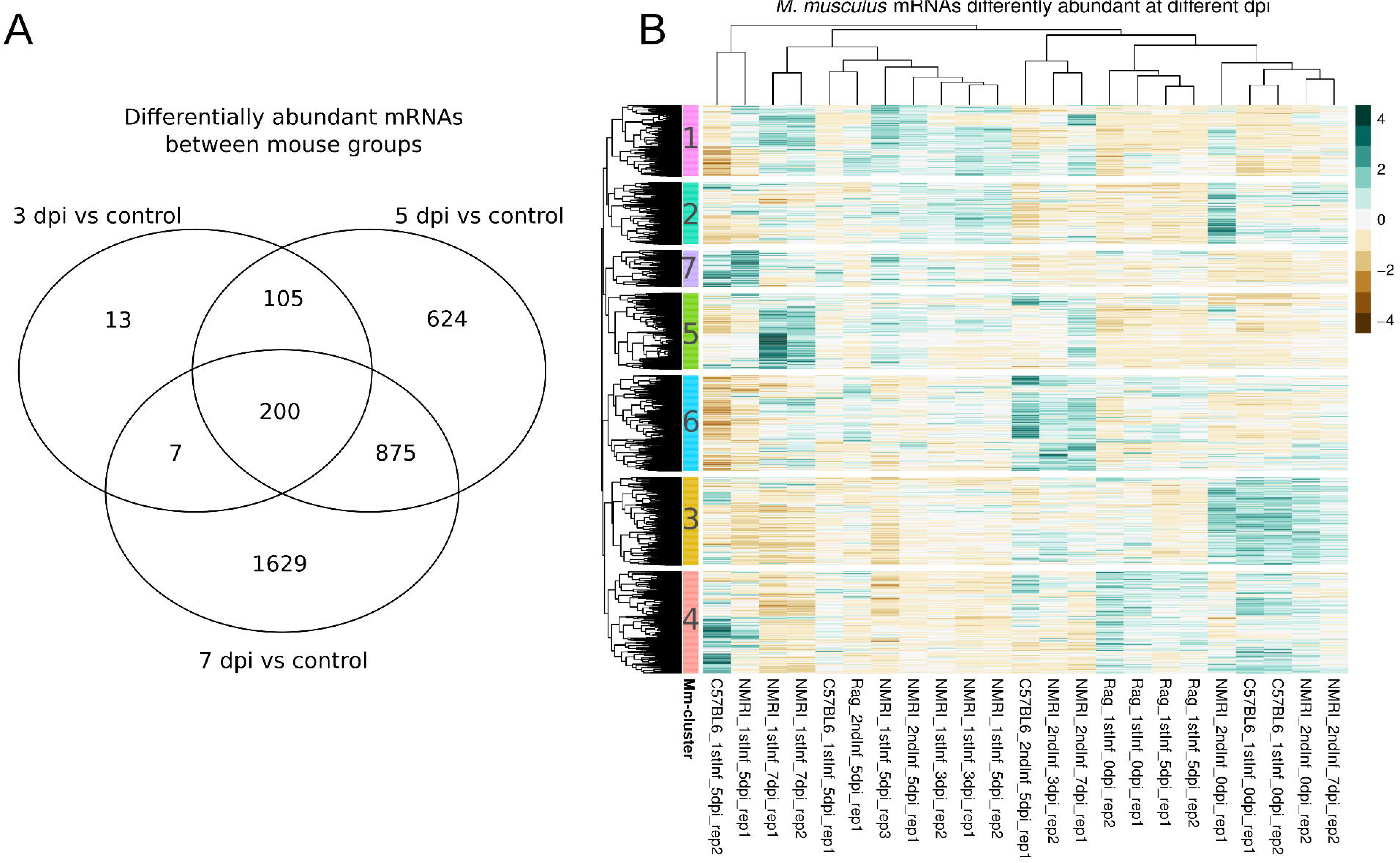
Differentially abundant mouse mRNAs and clustering thereof. A) Venn diagram visualizes the overlap between genes showing differential abundance (FDR < 0.01; edgeR glm likelihood-ratio tests) between i) uninfected controls and different time-points post infection and ii) between different time-points and the sum of all genes reacting to infection. Controls from challenge infection were used. B) Hierarchical clustering of differentially abundant mRNAs performed on Euclidean distances using complete linkage. Cluster cut-offs (dendrogram resolution) were set to identify gene-sets with profiles interpretable in relation to the parasite lifecycle and between mice of different immune competence.

To further analyze the distinct responses early and late in infection, we performed hierarchical clustering on transcript abundance patterns at different time-points post infection (Figure 2B). Three main sample clusters formed (dendrogram indicating similarities between columns at top of Figure 2B). Immune deficient *Rag1*^−/−^ mice, including infected *Rag1*^−/−^ samples, show an expression pattern most similar to uninfected samples. This similarity between infected and non-infected *Rag1*^−/−^ samples confirms the immune deficiency phenotype; a failure to react to infection in these mice, and suggests a strong influence of adaptive immune responses on overall transcriptional responses. Surprisingly, these patterns indicate that innate immune responses and other B- and T-cell independent processes play detectable though relatively small roles (mouse gene cluster 4; Mm-cluster hereafter, Figure 2B) in shaping the mouse transcriptome upon *E. falciformis* infection.

#### Responses to parasite infection differ between immunocompromised and immune competent mice

The self-limiting nature of *E. falciformis* infection and host resistance to reinfection ([33] and Figure 1A) makes it interesting to analyze transcriptomes of immune competent hosts in depth. On 3 and 5 days post infection, mRNAs of two clusters of genes have overall high abundance in samples of all immune competent infected animals (Mm-clusters 1 and 2). Other mRNAs (Mm-clusters 3 and 4) show lowered abundance in all those infected samples.

Gene Ontology (GO) terms enriched among the mRNAs which become more abundant only early in infection (Mm-clusters 1 and 2) are, e.g., “stem cell population maintenance”, “mRNA processing”, and “cell cycle G2/M transition”, indicating tissue remodeling in the epithelium. In addition, terms such as “regulation of response to food” are enriched (Table S1). This is interesting since weight losses and malnutrition are generally common during parasitic infections [34, 35], also in *Eimeria* spp. infections [36–38], and weight loss was also seen in the present study (Figure S4).

Genes whose mRNA levels decreased in abundance upon infection (Mm-clusters 3 and 4) indicate induction of IL-1 and IL-6, which are involved in inflammation, including T- and B-cell recruitment and maturation, and broad acute phase immune responses (Table S1). IL-6 has also been shown to support tissue repair and inhibit apoptosis after epithelial wounding [39]. In addition, IL-6 is linked to Th17 responses [40] which are known to play an important role in responses to *E. falciformis* [27]. Further terms indicate a regulation of transforming growth factor-β (TGFβ) which is important for wound healing in intestinal epithelium [41], epidermal growth factor (EGF) and tumor necrosis factor (TNF), which regulate proliferation of epithelial cells and inhibit apoptosis in epithelial cells [42,43]. Inhibition of Notch signaling, which is also highlighted by GO terms, has been shown to alter the composition of cell-types in the epithelium towards Paneth and Goblet-like cells [44].

Although speculative, several of the GO terms (e.g. “calcineurin-NFAT signaling cascade”, “Inositol-phosphate mediated signaling”, “Notch receptor processing” in addition to those mentioned above) annotated to genes whose mRNA levels change in abundance upon early infection (Mm-cluster 3 and 4) can be linked to explain fundamental mechanisms. Inositol signaling can lead to release of calcium and calcineurin-dependent translocation of NFAT to the nucleus; and there to activation of NFAT target genes in T-cells, but also many other cell types [45]. In addition, changes in the host epithelium do take place when cells are invaded by, e.g., *E. falciformis,* but also generally by pathogens, and this is reflected in the stem-cell and cell cycle-related GO terms described above for Mm-clusters 1 and 2. Further investigation of the role of the processes and molecules highlighted here will contribute to better understanding for epithelial responses to intestinal intracellular parasitic infection. Interestingly, in T- and B-cell deficient hosts, the same four groups of genes described above (Mm-clusters 1-4, Figure 2B), which are responsible for these dominating responses in immune competent hosts show no differences between infected and non-infected immune deficient animals.

#### Adaptive immune responses characterize late infection

Pronounced transcriptional changes in the mouse host occur late in infection in immune competent animals (Table 2 and Mm-cluster 5 in Figure 2B). Annotated processes and functions (GO terms) for genes with increased abundance at 7 days post infection reflect the expected onset of an adaptive immune response (Table S1). As late as 5 days post infection, genes responsible for these enrichments are still low on mRNA abundance. This confirms a strong induction of immune responses, particularly adaptive immune responses, between 5 and 7 days post infection. This result is well in line with previously described immune responses to infection with *Eimeria* spp. [23–27].

#### Protective responses occur earlier in challenge infected than in naïve hosts

Transcriptomes from three samples from early and late challenge infection show the same distinct profile of elevated mRNA abundance at 3, 5 and 7 days post infection (Mm-cluster 6, Figure 2B). The underlying mRNAs are highly enriched for GO terms for RNA processing, e.g., splicing, which indicated post-transcriptional regulation. In addition, terms for histone and chromatin modification are enriched (Table S1). This, along with less oocyst shedding during challenge infection, suggests that protective immune responses in challenge infected animals are regulated both at the transcriptional and post-transcriptional level. The high abundance of these mRNAs at different time-points post infection in wild-type hosts (NMRI) further indicates that protective immunity is similar at these time-points. Possibly, induction and chronologic differences in challenge infected animals occur before 3 days post infection. The completely cleared infection in some samples (Table 1; and unexpected clustering of e.g. NMRI_2ndInf_7dpi_rep2), apart from clearly demonstrating protection, also supports an early timing of this response upon challenge infection. However, the distinct shared profile at the investigated time-points (days 3, 5, and 7) does show that the protective response is still detectable at the transcriptional level several days after the challenge.

### A framework to interpret *E. falciformis* transcriptomes is provided by orthologues in the Coccidia *E. tenella* and *T. gondii*

To establish *E. falciformis* as a model for coccidian parasites, transcriptome profiles of orthologue genes from closely related parasites can help to draw parallels between lifecycle stages. This can be informative in predicting gene function and in analyzing evolutionary forces acting on the different lifecycle stages. Therefore, we performed correlation analysis between our *E. falciformis* transcriptome and RNA-seq transcriptomes from closely related parasites at corresponding stages of their lifecycles. Two datasets for the economically important chicken parasite *E. tenella* [46,47] and one dataset of the model apicomplexan parasite *T. gondii* [48] were included. The latter was used because it is to date the only available dataset for the complete in vivo lifecycle of *T. gondii* (including stages in the definitive cat host), and therefore compares well with our data.

For all samples from these studies and our data, abundances of orthologous genes were correlated and Spearman’s coefficient was compared (Figure 3). With the exception of sporozoites (see below), transcriptomes tend to be more strongly correlated (similar) between corresponding lifecycle stages of different parasite species than between stages in the same parasite species.

**Figure 3.**
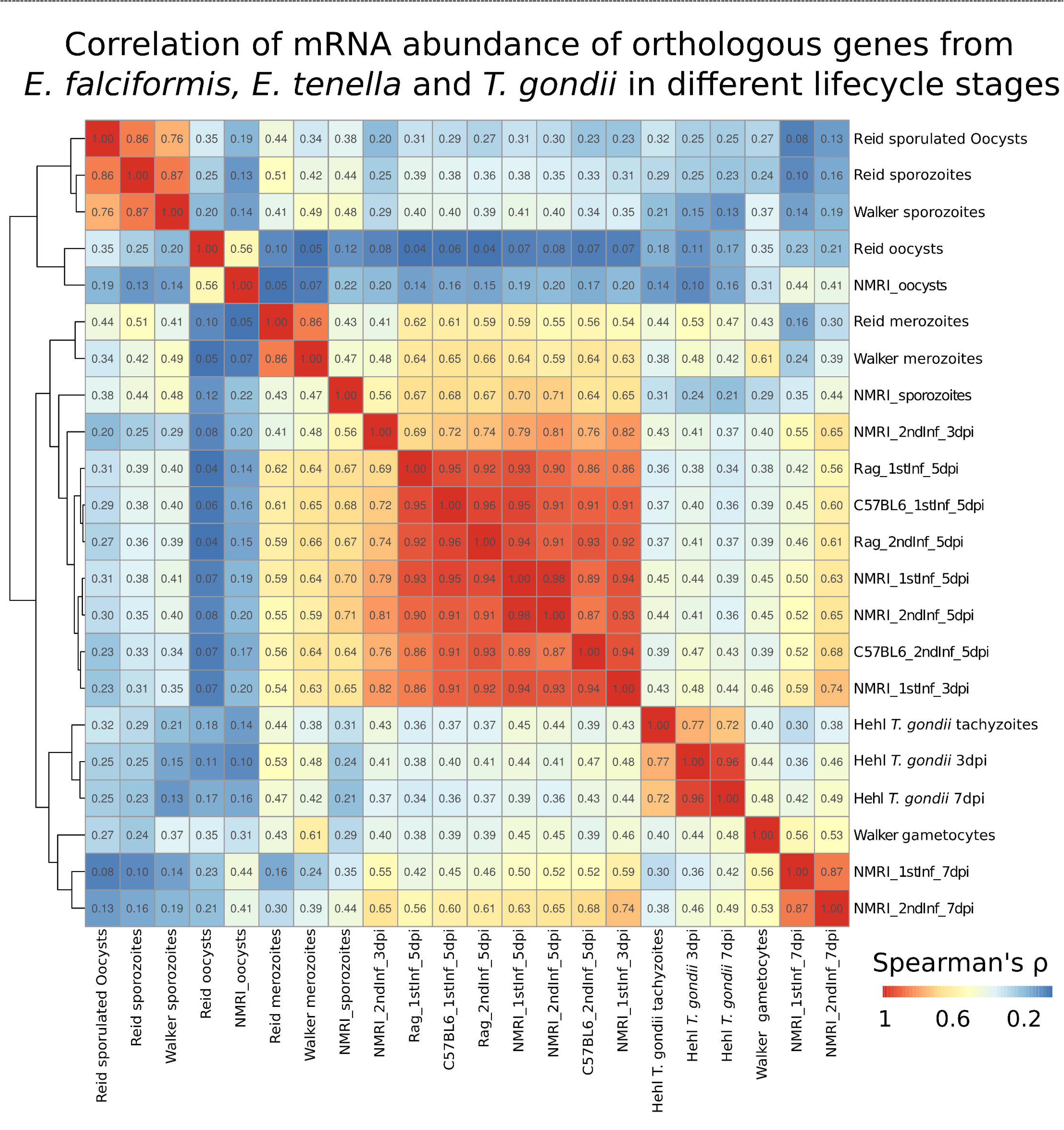
Correlations of *E. falciformis* mRNA abundance with ortholgues from other Coccidia. *E.falciformis* mRNA abundance was compared to that of orthologous genes of *E. tenella* [46,47] and *T. gondii* [48]. Correlation coefficients (Spearman’s ρ) were clustered using complete linkage. *T. gondii* and *Eimeria* spp. “late infection” samples cluster together. *E. falciformis* early infection samples cluster with *E. tenella* merozoites. *E. falciformis* sporozoites cluster with *E. falciformis* early infection, whereas unsporulated oocysts cluster with *E. tenella* unsporulated oocysts.

Orthologues in *E. tenella* and *E. falciformis* gamete stages (purified gametocytes and 7 dpi intestinal samples, respectively) are highly correlated in their expression across the two species, indicating conserved gene sets orchestrating sexual replication of the two parasites. Similarly, transcriptomes of *E. tenella* merozoites from both independent studies of that parasite are most similar to early *E. falciformis* samples, indicating similarity also during asexual reproduction. *E. falciformis* unsporulated oocyst transcriptomes share the highest similarity with those of unsporulated *E. tenella* oocysts.

*E. falciformis* sporozoites transcriptome profiles are more similar to *E. falciformis* early infection samples than to sporozoite transcriptomes of *E. tenella* orthologues. Similarities between sporozoites and early infection stages could be explained by similar biological processes, especially host cell invasion (and reinvasion by merozoites), being prepared or performed. Sporozoites are the only lifecycle stages in which orthologue mRNA abundance patterns show such dissimilarities to *E. tenella* and this might indicate a higher species specificity of the genes and processes in this invasive stage. This could be a result of virulence factors being expressed in this stage, which are known to undergo rapid gene family expansion, as seen in SAGs in *E. falciformis* [16], *T. gondii* [49], *Neospora caninum* [50], and other *Eimeria* spp. [46], or *var* genes in *Plasmodium falciparum* [51].

Below we provide a detailed description of the *E. falciformis* transcriptome, including a discussion of genes which have been shown to be important in closely related parasites such as *E. tenella* and *T. gondii.*

#### *Overall transcriptional changes in the lifecycle of* E. falciformis

Similar to the host transcriptome, differences in parasite mRNA abundance were mostly observed between late and early infection. Between 3 and 5 dpi 103 mRNAs were differently abundant (edgeR likelihood ratio tests on glms; FDR < 0.01), whereas between 3 and 7 dpi 1399 mRNAs, and between 5 and 7 dpi 2084 mRNAs were differentially abundant (Figure 4A). Hierarchical clustering did not group samples from 3 and 5 days distinctively and we thus refer to these as “early infection” and 7 dpi as “late infection”. Distinct abundance differences define early infection (parasite gene cluster 6, “Ef-cluster” hereafter, Figure 4B). At those time-points asexual reproduction takes place [17,19]. Two separate clusters define late infection (7 dpi, Ef-clusters 2 and 7) in which we assume gametocytes to be present due to the peak of oocyst shedding one day later (Figure 1A) [17] and similarity of these transcriptomes with purified *E. tenella* gametocytes (Figure 3). The extracellular stages, sporozoites (Ef-cluster 4) and unsporulated oocysts (Ef-clusters 1 and 5) are clearly distinct by high mRNA abundance. In order to assess the biological relevance of these patterns, we applied enrichment analyses for GO terms and “gene family conservation profiles” based on earlier annotations [16].

**Figure 4.**
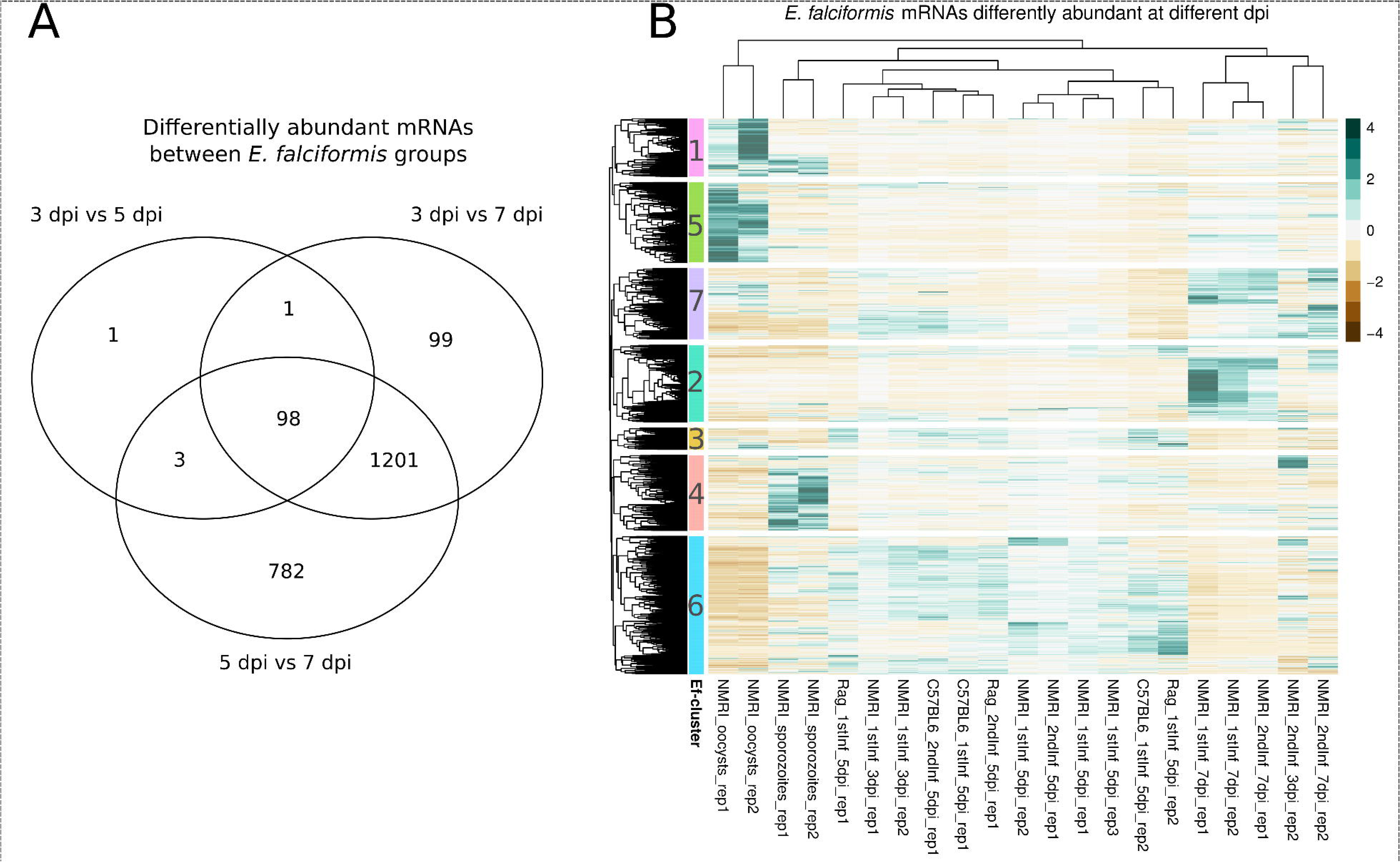
Differentially abundant *E. falciformis* mRNAs and clustering thereof. A) Venn diagram visualizes the overlap between genes showing differential abundance (FDR < 0.01; edgeR glm likelihood-ratio tests) between intracellular stages at 3 days post infection, 5 days post infection and 7 days post infection. B) Hierarchical clustering of abundance profiles for differentially abundant mRNAs performed on Euclidean distances using complete linkage. Cluster cut-offs (dendrogram resolution) were set to identify gene-sets with profiles interpretable in relation to the parasite lifecycle.

#### *Sporozoites express genes which are evolutionarily unique to* E. falciformis

Sporozoites are in our study released from oocysts in vitro, after which they are capable of invading host cells. We suggest that the requirement for proteins which mediate motility and other invasion processes are reflected by their mRNA levels in the transcriptome. We find that *E. falciformis* sporozoites are defined by a group of genes (Ef-cluster 4, Figure 4B) that is largely specific to *E. falciformis* (Table 3). This indicates that *E. falciformis* does not share with other species many of the abundant sporozoite genes so far described for those Coccidia. Interestingly, five out of 12 SAG gene transcripts predicted for *E. falciformis* [16] are typical for sporozoites. SAG proteins are thought to be involved in host cell attachment and invasion, and possibly in induction of immune responses in other apicomplexan species [46,50,52–56]. In total, mRNAs encoding ten SAGs were detected as differentially abundant in our data, but in other lifecycle stages than sporozoites. Such expression of particular SAGs in stages other than sporozoites has been reported for *E. tenella* [57]. Genes also receiving attention as potential virulence factors in *E. tenella* are rhoptry kinases (RopKs) [58]. Transcripts of two out of ten *E. falciformis* orthologues of RopKs are highly abundant in sporozoites (Ef_cluster 4). Also in *E. tenella* some RopKs are expressed predominantly in sporozoites and have been shown to be differentially expressed compared to *E. tenella* intracellular merozoite stages [59]. For genes with orthologues known to be important in other Coccidia, e.g., SAGs and RopKs, orthologues indicate a molecular function, but the biological relevance of their expression in *E. falciformis* remains unclear.

**Table 3.** Enrichments and underrepresentation of species or species-group orthologues in *E. falciformis* gene clusters (from Figure 3b). Odds ratios higher than one indicate enrichment and smaller than one indicate underrepresentation. Conservation categories were chosen as previously described [16]. Only significant results (FDR < 0.05) are shown.

| *E. falciformis* cluster | Conservation category | Odds ratio | p-value | FDR |
| --- | --- | --- | --- | --- |
| Ef-cluster 2 (up at 7 dpi) | Conserved | 0.67 | 9.03E-06 | 1.90E-04 |
| Ef-cluster 4 (up in sporozoites) | Conserved | 0.72 | 2.44E-04 | 1.71E-03 |
| Ef-cluster 7 (up at 7 dpi) | Conserved | 1.72 | 1.11E-10 | 4.65E-09 |
| Ef-cluster 2 (up at 7 dpi) | ApicomplexaC | 0.45 | 1.84E-04 | 1.71E-03 |
| Ef-cluster 5 (up in oocysts) | ApicomplexaC | 1.86 | 3.76E-05 | 5.26E-04 |
| Ef-cluster 4 (up in sporozoites) | *E. falciformis* | 3.05 | 2.38E-04 | 1.71E-03 |
| Ef-cluster 1 (up in oocysts) | *Eimeria* | 0.68 | 1.83E-03 | 9.59E-03 |
| Ef-cluster 6 (up in early inf) | Apicomplexa | 1.46 | 1.11E-03 | 6.64E-03 |

Genes typical for the sporozoite stage displayed a species specific profile with the respective gene families absent outside *E. falciformis* (Table 3). This mirrors our analysis of orthologous genes, in which sporozoites were the only lifecycle stage not displaying strong cross-species correlation in their transcriptome. This suggests that traits involved in host cell invasion may have evolved quickly and rapidly become specific for a parasite in its respective host species or target organ niche.

For the overall biological functions of sporozoite genes (Ef-cluster 4), GO enrichment data suggests ATP production and biosynthesis processes as dominant features (Table S2). In addition, this invasive stage is characterized by “maintenance of protein location in cell” and GO terms which indicate similar biological functions. Possibly, this reflects control of microneme or rhoptry protein localization as sporozoites prepare for invasion. Sporozoites therefore display a transcriptome indicative of large requirements for ATP and production of known virulence factors such as SAG and RopKs and are characterized by expression of species specific genes.

Genes typical for the sporozoite stage displayed a species specific profile with the respective gene families absent outside *E. falciformis* (Table 3). This mirrors our analysis of orthologous genes, in which sporozoites were the only lifecycle stage not displaying strong cross-species correlation in their transcriptome. This suggests that traits involved in host cell invasion may have evolved quickly and rapidly become specific for a parasite in its respective host species or target organ niche.

#### Growth processes dominate the transcriptome during asexual reproduction

Invasion of epithelial cells by sporozoites is followed by asexual reproduction leading to a massive increase in parasite numbers between 3 and 5 days post infection, when several rounds of schizogony take place in a somewhat unsynchronized fashion [17,19]. In early infection, and similar to sporozoites, mRNAs annotated for biosynthetic activity are enriched, but different genes/mRNAs are contributing to enrichment of similar GO terms compared to sporozoites (Table S2). Enrichment of terms referring to replication and growth-related processes (biosynthesis) highlights the parasite’s expansion during schizogony.

Amongst early infection high abundance mRNAs, we found four out of ten RopKs which are predicted in *E. falciformis* [16]. This is the largest number of RopKs in any one group of differentially abundant mRNAs in our analysis and they constitute a statistically significant enrichment (Fisher’s exact test; p < 0.001). Three of these have orthologues in *T. gondii*: ROP41, ROP35 and ROP21 [60–63]. Our data gives a first overview of expression patterns for *E. falciformis* RopKs and offer a good starting point for functional analysis of these virulence factors in *Eimeria* spp..

#### Gametocyte motility dominates the transcriptome late in infection

Two *E. falciformis* gene clusters show a distinct profile characterized by high mRNA abundance on 7 days post infection (Ef-clusters 2 and 7; Figure 4B). Both clusters display low mRNA abundance in other lifecycle stages, especially in oocysts and sporozoites. Enriched GO terms such as “movement of cell or subcellular component” and “microtubule-based movement” along with terms suggesting ATP production (e.g. “ATP generation from ADP”) indicate the presence of motile and energy demanding gametocytes in these samples. Peptide and nitrogen compound biosynthetic processes along with “chitin metabolic process” (Table S2) also suggest that the parasite produces building blocks for oocysts and their walls in this stage. Our data confirms findings of Walker et al. (2015) in *E. tenella* gametocytes: these authors also identified cytoskeleton related and transport processes as upregulated in gametocytes compared to merozoites or sporozoites [47].

#### Oocysts are characterized by cell differentiation and DNA replication processes

Oocysts are the infective stage in the lifecycle of Coccidia. They are shed with feces as unsporulated, “immature”, capsules and in the environment they undergo sporulation - meiotic and mitotic divisions [14] – and become infective. Our oocysts were purified in the unsporulated stage from passage in lab mice. Two expression clusters of mRNA are highly abundant in this stage (Ef-clusters 1 and 5; Figure 4B). One of these oocyst gene sets (Ef-cluster 5) is enriched for apicomplexan-shared orthologues (Table 3) and for GO terms such as “DNA repair”, “protein modification process” and “cell differentiation”, supporting that expected sporulation processes have been initiated. The same cluster is also the only cluster which is enriched for transmembrane domains (Fisher’s exact test, FDR < 0.001).

#### *E. falciformis* does not respond plastically to differences in the host transcriptome

We show that infections of *E. falciformis* in its natural host, the house mouse, follow a genetically canalized and chronological pattern independent of the immune status of the host. This is supported by the lack of separation of parasite transcriptomes from immune competent and immune deficient hosts, or from naïve and challenge infected hosts (Figure 4B). In the immune competent host, a switch from epithelial remodeling and innate immune processes to adaptive immune responses between 5 and 7 days post infection are paralleled by a parasite switch from asexual to sexual reproduction. This contemporaneity might be an evolutionary adaptation of the parasite to host responses in order to finish its lifecycle before the host environment becomes hostile. Such a response could be based on a) genetically canalized developmental timing or b) the parasite sensing an immune challenge and establishing a reaction, i.e. respond plastically. However, in an immune deficient host, which lacks the described responses in its transcriptome, the parasite’s transcriptome cannot be distinguished from one in an immune competent host. We thereby provide evidence from hosts with variation in their immune responses that support that *E. falciformis* follows a non-plastic, and instead genetically canalized program during its lifecycle in the mouse host.

### Conclusion

In this dual transcriptome study, we provide a thorough description of transcriptional responses in mice to infection with *E. falciformis,* and corresponding parasite transcriptomes. The mouse epithelial transcriptome of naïve, immune competent mice changes upon infection. Responses in wild-type challenge infected hosts suggest strong regulation both at the transcriptional level and in RNA processing. In contrast, these patterns are missing in immunocompromised animals which instead show a minimal transcriptional response to infection, demonstrating the host dependence of mature T- and B-cells for a natural response to this coccidian parasite.

For the first time we also describe the full parasite lifecycle transcriptomes of *E. falciformis.* Parasite transcriptomes are not distinguishable between hosts of different immune competence, demonstrating lack of plasticity at the gene expression and mRNA levels. Two independent assessments of evolutionary conservation show that invasive sporozoites possess the most species-specific transcriptomes in the *E. falciformis* lifecycle. We therefore suggest that excysted sporozoites express most of the genes involved in host-parasite coevolutionary processes, which accelerate divergence and may determine niche specificity.

Taken together, we propose that *E. falciformis* follows a genetically predetermined path rather than responding to cues from the host, such as differences in immune responses.

We further suggest that analyzing plasticity in parasites and comparing this between different host genotypes or species can be a useful tool to understand the evolutionary development of niche specificity or a generalist parasitic life-style infecting multiple different hosts or tissues. We emphasize that gene expression is not necessarily a product of plastic host-parasite interactions, especially not in the parasite, but may instead follow genetically determined programs.

## METHODS

### Mice, infection procedure and infection analysis

Three strains of mice were used in our experiments: NMRI, C57BL/6 (Charles River Laboratories, Sulzfeld, Germany), and *Rag1*^−/−^ on C57BL/6 background (obtained from German Rheumatism Research Centre, Berlin). *Rag1*^−/−^-mice are deficient in T- and B-cell maturation. Animals where infected as described by Schmid et al. [64], but tap-water was used instead of PBS for administration of oocysts. Briefly, NMRI mice were infected two times, which will be referred to as naïve and challenge infection. For the naïve infection, 150 sporulated oocysts were administered in 100 μL water by oral gavage. During the naïve infection of 52 mice, all animals were weighed every day. On day zero, before infection, as well as on 3 dpi, 5 dpi and 7 dpi, ceca from 3-4 sacrificed mice per time-point were collected. Epithelial cells were isolated as described in Schmid et al. (2012), in which the protocol generated epithelial cells with 90 *%* purity. For challenge infection, mice recovered spontaneously and were after four weeks challenge infected. Recovery was monitored by weighing and visual inspection of fur. For the challenge infection, 1500 sporulated oocysts were applied by oral gavage in 100μL water (a higher dose was necessary to establish a challenge infection). Tissue from three to four mice per replicate was pooled for both non-reinfection control (referred to as day 0 of challenge infection) and for all other samples. *Rag1*^−/−^ mice and the background C57BL/6 strain control mice were also subjected to naïve and challenge infections with 10 sporulated oocysts in 100 μL water in both cases. Samples were taken on day 0 (pre-infection control) and 5 dpi in both naïve and challenge infections of these mice and were otherwise treated as described above for NMRI mice. Oocyst shedding was determined from eight NMRI mice in naïve infection and four challenge infected, from 15 naïve Rag1^−/−^ and C57BL/6 mice respectively, and from nine challenge infected Rag1^−/−^ and C57BL/6 mice, respectively. Overall oocyst output was compared using Mann-Whitney U-test in R [65].

### Oocyst purification for infection, sequencing and quantification

Oocysts for infection were purified by NaOCl flotation of mouse feces stored in potassium dichromate, in which oocysts for infection were allowed to sporulate at room temperature for at least five days. During the patency phase, feces of mice were collected and oocysts were flotated using saturated NaCl-solution. The oocyst output was quantified using the McMaster chamber. For sequencing, unsporulated oocysts were purified twice per day from feces of NMRI mice on 8 – 10 dpi, and immediately subjected to RNA purification. The strain “*E. falciformis* Bayer Haberkorn 1970” was used for all infections and parasite samples, it is maintained through passage in NMRI mice in our facilities as described previously [64].

### Sporozoite isolation

Sporocysts were isolated according to the method of [66] with slight modifications. Briefly, not more than 5 million sporulated oocysts were resuspended in 0.4% pepsin solution (Applichem), pH 3, and incubated at 37°C for 1 hour. Subsequently, sporocysts were isolated by mechanical shearing using glass beads (diameter 0.5 mm), washed and separated from oocyst cell wall components by centrifugation at 1800 g for 10 min. Sporozoites were isolated from sporocysts by in vitro excystation. For this, sporocysts were incubated at 37°C in DMEM containing 0.04% tauroglycocholate (MP Biomedicals) and 0.25% trypsin (Applichem) for 30 min. Released sporozoites were purified in cellulose columns as described in [67].

### RNA extraction and quantification

For RNA-seq, total RNA was isolated either from infected epithelial cells, sporozoites, or unsporulated oocysts using Trizol according to the manufacturer’s protocol (Invitrogen). In addition, unsporulated oocysts in Trizol were treated by mechanical shearing using glass beads for at least 20 min under frequent microscopic inspection. Purified RNA was used to produce an mRNA library using Illumina’s TruSeq RNA Sample Preparation guide. For qPCR, uninfected and infected epithelial cells from 3, 5 and 7 dpi were isolated as described above and stored in 1 mL Trizol. Total RNA was isolated using the PureLink RNA Mini Kit (Invitrogen) and reverse transcribed into cDNA using the Superscript III Platinum Two Step qRT-PCR Kit (Thermo Fisher Scientfic).

These RNA preparations were used for RT-qPCR of *Eimeria* 18S and creation of a mouse gene reference index. For the reference index, the mouse genes cytochrome c-1 (Cyc), peptidylprolyl isomerase A (Ppia) and peptidylprolyl isomerase B (Ppib) were amplified using the primers Cyc1_qPCR_f (5’-CAGCTACCATGTCACAAGTAGC-3’) and Cyc1_qPCR_r (5’- ACCACTTATGCCGCTTCATG −3’); Ppib_qPCR_f (CAAAGACACCAATGGCTCAC) and Ppib_ qPCR_r (5’-TGACATCCTTCAGTGGCTTG-3’); Ppia_qPCR_f (5’-ACCGTGTTCTTCGACATCAC-3’) and Ppia_qPCR_r (5’-ATGGCGTGTAAAGTCACCAC-3’), respectively. The *E. falciformis* 18S gene was amplified using the primers Ef18s_for (5’-ACAATTGGAGGGCAAGTCTG-3’) and Ef18s_rev (5’-AAACACCAACAGACGCAGTG-3’).

After initialization at 50°C followed by activation of enzymes at 95°C, 40 amplification cycles consisting of denaturation at 95°C for 15s and combined annealing and elongation at 60°C for 60s were performed. After each cycle the fluorescent signal was measured. A reference index was constructed taking the cube route of the multiplied crossing threshold (ct)-values for the tree mouse genes. This composite “index ct-value” was used to calculate the ct difference (delta-ct) of the *E. falciformis* 18S gene. The procedure was performed in technical triplicate for each sample and mean delta-ct values were taken. A linear model was constructed in R [65] to predict these normalized delta-ct values by day post infection (dpi) and type of infection (naïve or challenge infected). This model excludes measurements at 0 days post infection as background noise.

### Sequencing and quality assessment

cDNA libraries were sequenced on either GAIIX (13 samples) or Illumina Hiseq 2000 (14 samples) platforms in a total of four batches (different machine runs) as specified in Table 1. A fastq_quality_filter (FASTQ-toolkit, version 0.0.14, available at https://github.com/agordon/fastx_toolkit.git) was applied to Illumina Hiseq 2000 samples using a phred score of 10. We intentionally did not use a stringent trimming before mapping to genome assemblies as the mapping process itself has been shown to be a superior quality control [68].

### Alignment and reference genomes

The *Mus musculus* mm10 assembly (Genome Reference Consortium Mouse Build 38, GCA_000001635.2) was used as reference genome for mapping and corresponding annotations were used for downstream analyses. The *E. falciformis* genome [16] was downloaded from ToxoDB [49]. For mapping, mouse and parasite genome files were merged into a combined reference genome, and files including mRNA sequences from both species were aligned against this reference using TopHat2, version 2.0.14, [69] with the option-G specified, and Bowtie2, version 1.1.2, [70]. This was done to avoid spurious mapping in ultraconserved genomic regions. Single-end and pair-end sequence samples were aligned separately with library type ‘fr-unstranded’ specified for pair-end samples. Bam files were used as input for the function “featureCounts” from of the R package “Rsubread” [71]. All subsequent analyses were performed in R [65].

### Differential mRNA abundance, data normalization and sample exclusions

After import of data to R, mouse and parasite data was separated using transcript IDs and analyzed, including normalization, separately. For each species, count data was normalized using the R-package edgeR version 3.16.2 [72] with the upperquartile normalization method.

This raw data underlying our study is available as supplementary data S1. Briefly, genes with below an overall of 3000 reads (mouse) and 100 reads (*E. falciformis*) summed over all samples (libraries) were removed and normalization factors were calculated for the 75% quantile for each library. This normalization is suitable for densities of mapping read counts which follow a negative binomial distribution. Technically, this exclusion made it possible to obtain parasite read counts in agreement with a negative binomial distribution. We excluded samples NMRI_2nd_3dpi _rep1 and NMRI_2nd_5dpi_rep2 due to low parasite contribution (0.012% and 0.023%) to the overall transcriptome. Technically, this exclusion made it possible to obtain parasite read counts in agreement with a negative binomial distribution. Both excluded samples are from challenge infection and it is likely that the infected mice were immune to re-infection. One additional sample (NMRI_1stInf_0dpi_rep1) was excluded because the uninfected control showed unexpected mapping of reads to the *E. falciformis* genome (0.033%). As samples and individual replicates were sequenced in batches to different depth and using different instrumentation (Table 1) we performed multidimensional scaling of samples as quality controls using the function “plotMDS” provided in the R package edgeR v 3.16.2 [72].

### Testing of differentially abundant mRNAs and hierarchical clustering

We used edgeR v 3.16.2 [72] further to fit generalized linear models (GLMs with a negative binomial link function) for each gene (glmFit) and to perform likelihood ratio tests for models with or without a focal factor (glmLRT) using the “alternate design matrix” approach specifying focal contrasts individually. Tested contrasts comprised for the mouse a) infections at each time-point versus uninfected controls, b) corresponding time-points between different mouse strains and c) corresponding time-points and mouse strains for naïve and challenge infection. Since the control sample for infection in naïve NMRI mice was removed from the analysis (see above), the two uninfected replicates from challenge infection were used as uninfected controls in all NMRI mouse analyses. For the parasite, contrasts were set between a) all different stages of the lifecycle, as well as b) and c) as above (see also results in Table 2).

Mouse mRNAs which responded to infection or were differently abundant at different time-points of infection (0 vs “any days post infection” or “any days post infection” vs “any days post infection”; see Table 2) and *E. falciformis* genes showing differences between any lifecycle stage (oocysts versus sporozoites, or either of those versus “any days post infection” or “any days post infection” versus “any days post infection”) were selected and used for hierarchical clustering. Hierarchical clustering was performed using the complete linkage method based on Euclidean distances between Z-scores (mRNA abundance values scaled for differences from mean over all samples of each gene in units of standard deviations).

### Enrichment tests and evolutionary conservation test

Gene Ontology (GO) enrichment analysis was performed using the R package topGO with the “weight0l” algorithm and Fisher’s exact tests. We additionally performed a correction for multiple testing on the returned p-values (function “p.adjust” using the BH-method [73]). Similarly, a Fisher’s exact test and corrections for multiple testing were used to test for overrepresentation of transcripts with a signal sequence for entering the secretory pathway or containing transmembrane domains (as inferred using Signal P) which are predicted for the *E. falciformis* genome [16]. Evolutionary conservation of gene families was analyzed based on categories from [16] which are as follows: i) *E. falciformis* specific, ii) specific to the genus *Eimeria,* compiled by an analysis of *E. falciformis, E. maxima* and *E. tenella,* iii) Coccidia: *Eimeria* plus *T. gondii* and *Neospora caninum,* iv) Coccidia plus *Babesia microti, Theileria annulata, Plasmodium falciparum* and *Plasmodium vivax* v) the same apicomplexan parasites as in iv plus *Cryprosporidium hominis,* vi) universally conserved in the eukaryote superkingdom inferred from an analysis of *Saccharomyces cerevisiae* and *Arabidopsis thaliana.* These categories were tested for overrepresentation in parasite gene clusters with particular patterns described in the text using Fisher’s exact-tests. Resulting p-values were corrected for multiple testing using the procedure of Benjamini and Hochberg [72] and reported as false discovery rates (FDR).

### Correlation analysis of apicomplexan transcriptomes

Transcriptome datasets from [46,47] and [48] were downloaded from ToxoDB [49].

Orthologues between *E. falciformis, E. tenella* and *T. gondii* were compiled as in [16] and only 1:1:1 orthologue triplets were retained for analysis, as multi-paralog gene-families might contain members showing divergent evolution of gene-expression due to neo/sub functionalization. Mean mRNA abundances per lifecycle stage were used for samples from our study. Spearman’s correlation coefficients for expression over different samples in all studies and over different species represented by their orthologues were determined. Hierarchical clustering with complete linkage was used to cluster resulting correlations coefficients.

## DECLARATIONS

### Ethics approval and consent to participate

Animal procedures were performed according to the German Animal Protection Laws as directed and approved by the overseeing authority Landesamt fuer Gesundheit und Soziales (Berlin, Germany) under numbers H0098/04 and G0039/11.

### Consent to publish

Not applicable

### Availability of data and materials

Raw data will be deposited to ENA/SRA. A processed version of this data will be available at ToxoDB (http://toxodb.org/toxo/) for interactive analysis and download. Code underlying our analysis and intermediate result files are available at https://github.com/derele/Ef_RNAseqtagged as version 0.1. And will be deposited upon acceptance in it's final version at Zenodo (https://zenodo.org/).

### Authors' contributions

TE, SS, CD, RL and EH designed the experiments, RL performed infections, EH obtained grant support for the work, RL, SS, CD and EH gathered the data, EH and TE analyzed the data, TE and EH drafted the manuscript, TE, SS, RL and EH edited the manuscript, all authors contributed original ideas to the research and agreed on the final version of the manuscript.

## ACKNOWLEDGEMENTS

The authors wish to thank Frank Seeber and Toni Aebischer for valuable comments on the manuscript, Annica Rebbig for establishing the mouse reference index for RT-qPCR and Kirsi Blank for support in oocyst purification and counting, parasite passaging and RT-qPCRs.

## SUPPLEMENTARY INFORMATION

### Supplementary Figures

**Figure S1.** Ordinations on mouse and parasite transcriptomes. The results of multidimensional scaling analyses are displayed for mouse and *E. falciformis* using different labels to allow comparisons.

**Figure S2.** Controls for the properties of mRNA abundance distributions after setting different abundance thresholds per mRNA over all samples.

**Figure S3.** Mouse mRNA abundance in late *E. falciformis* infection versus uninfected controls, assessed by both RNA-seq (present data) and microarray. Mouse data from 7 days post infection (RNA-seq) and 6 days post infection. In both experiments, NMRI mice were infected with the same E. falciformis isolate. Even with one day difference in sampling, mouse transcriptomes show a strong correlation. The line depicted for visualisation corresponds to generalized additive model unsing penalized regression splines.

**Figure S4.** Weight loss of mice during *E. falciformis* infection.

Mouse weight is shown as a percentage relative to weight at the time of infection. Infection dose for NMRI was 150 oocysts in naïve infection and 1500 in challenge infection. For C57BL/6 and Rag1-/-dose was 10 oocysts in both naïve and challenge infection. Bars indicate standard error for three or four replicates.

### Supplementary Tables

Table S1. GO terms enriched in Mm-clusters in Figure 2B.

Table S2. GO terms enriched in Ef-clusters in Figure 4B.

## REFERENCES

1. Stearns SC. The evolutionary significance of phenotypic plasticity. BioScience. 1989;39:436–45.

2. Dodson S. Predator-induced reaction norms. BioScience. 1989;39:447–52.

3. Pancer Z, Cooper MD. The evolution of adaptive immunity. Annu. Rev. Immunol. 2006;24:497–518.

4. Viney M, Diaz A. Phenotypic plasticity in nematodes. Worm. 2012;1:98–106.

5. Stear MJ, Bairden K, Duncan JL, Holmes PH, McKellar QA, Park M, et al. How hosts control worms. Nature. 1997;389:27–27.

6. Weclawski U, Heitlinger EG, Baust T, Klar B, Petney T, Han Y-S, et al. Rapid evolution of Anguillicola crassus in Europe: species diagnostic traits are plastic and evolutionarily labile. Front. Zool. 2014;11:74.

7. Weclawski U, Heitlinger EG, Baust T, Klar B, Petney T, San Han Y, et al. Evolutionary divergence of the swim bladder nematode Anguillicola crassus after colonization of a novel host, Anguilla anguilla. BMC Evol. Biol. 2013;13:78.

8. Heitlinger E, Taraschewski H, Weclawski U, Gharbi K, Blaxter M. Transcriptome analyses of Anguillicola crassus from native and novel hosts. PeerJ. 2014;2:e684.

9. Weber C, Koutero M, Dillies M-A, Varet H, Lopez-Camarillo C, Coppée JY, et al. Extensive transcriptome analysis correlates the plasticity of Entamoeba histolytica pathogenesis to rapid phenotype changes depending on the environment. Sci. Rep. 2016. http://www.ncbi.nlm.nih.gov/pmc/articles/PMC5073345/. Accessed 23 Feb 2017.

10. Lovegrove FE, Peña-Castillo L, Mohammad N, Liles WC, Hughes TR, Kain KC. Simultaneous host and parasite expression profiling identifies tissue-specific transcriptional programs associated with susceptibility or resistance to experimental cerebral malaria. BMC Genomics. 2006;7:295.

11. da Silva F, da Fonseca M, Barbosa HS, Gross U, Lüder CGK. Stress-related and spontaneous stage differentiation of Toxoplasma gondii. Mol. Biosyst. 2008;4:824–34.

12. Buchholz KR, Fritz HM, Chen X, Durbin-Johnson B, Rocke DM, Ferguson DJ, et al. Identification of tissue cyst wall components by transcriptome analysis of in vivo and in vitro Toxoplasma gondii bradyzoites. Eukaryot. Cell. 2011;10:1637

13. Sibley DL, Charron A, Håkansson S, Mordue D. Invasion and Intracellular Survival by Toxoplasma. In: Madame Curie Bioscience Database [Internet]. Austin (TX): Landes Bioscience; 2000-2013. Available from: https://www.ncbi.nlm.nih.gov/books/NBK6450/2013.

14. Duszynski DW. Eimeria, Eimeria. eLS. John Wiley & Sons, Ltd. 2011. http://onlinelibrary.wiley.com/doi/10.1002/9780470015902.a0001962.pub2/abstract. Accessed 10 Oct 2016.

15. Chapman HD, Barta JR, Blake D, Gruber A, Jenkins M, Smith NC, et al. A selective review of advances in coccidiosis research. Adv. Parasitol. 2013;83:93.

16. Heitlinger E, Spork S, Lucius R, Dieterich C. The genome of Eimeria falciformis - reduction and specialization in a single host apicomplexan parasite. BMC Genomics. 2014;15:696.

17. Haberkorn A. Die Entwicklung von Eimeria falciformis (Eimer 1870) in der weißen Maus (Mus musculus). Z. Für Parasitenkd. 1970;34:49–67.

18. Schmid M, Heitlinger E, Spork S, Mollenkopf H-J, Lucius R, Gupta N. Eimeria falciformis infection of the mouse cecum identifies opposing roles of IFNγ-regulated host pathways for the parasite development. Mucosal Immunol. 2013;7:969–982

19. Mesfin GM, Bellamy JEC. Effects of acquired resistance on infection with Eimeria falciformis var. pragensis in mice. Infect. Immun. 1979;23:108–14.

20. Montes C, Rojo F, Hidalgo R, Ferre I, Badiola C. Selection and development of a Spanish precocious strain of Eimeria necatrix. Vet. Parasitol. 1998;78:169–83.

21. Pakandl M. Selection of a precocious line of the rabbit coccidium Eimeria flavescens Marotel and Guilhon (1941) and characterisation of its endogenous cycle. Parasitol. Res. 2005;97:150–5.

22. Rose ME. Immune responses in infections with Coccidia: Macrophage activity. Infect. Immun. 1974;10:862–71.

23. Blagburn BL, Todd KS. Pathological changes and immunity associated with experimental Eimeria vermiformis infections in Mus musculus. J. Protozool. 1984;31:556–61.

24. Rose ME, Hesketh P, Wakelin D. Immune control of murine coccidiosis: CD4+ and CD8+ T lymphocytes contribute differentially in resistance to primary and secondary infections. Parasitology. 1992;105:349–54.

25. Gadde U, Chapman HD, Rathinam TR, Erf GF. Acquisition of immunity to the protozoan parasite Eimeria adenoeides in turkey poults and the peripheral blood leukocyte response to a primary infection. Poult. Sci. 2009;88:2346–52.

26. Sühwold A, Hermosilla C, Seeger T, Zahner H, Taubert A. T-cell reactions of Eimeria bovis primary-and challenge-infected calves. Parasitol. Res. 2010;106:595–605.

27. Stange J, Hepworth MR, Rausch S, Zajic L, Kühl AA, Uyttenhove C, et al. IL-22 mediates host defense against an intestinal intracellular parasite in the absence of IFN-γ at the cost of Th17-driven immunopathology. J. Immunol. Baltim. Md 1950. 2012;188:2410–8.

28. Foth BJ, Zhang N, Chaal BK, Sze SK, Preiser PR, Bozdech Z. Quantitative time-course profiling of parasite and host cell proteins in the human malaria parasite Plasmodium falciparum. Mol. Cell. Proteomics MCP. 2011;10:1–16.

29. Li Y, Shah-Simpson S, Okrah K, Belew AT, Choi J, Caradonna KL, et al. Transcriptome remodeling in Trypanosoma cruzi and human cells during intracellular infection. PloS Pathog. 2016;12:e1005511.

30. Fernandes MC, Dillon LAL, Belew AT, Bravo HC, Mosser DM, El-Sayed NM. Dual transcriptome profiling of Leishmania-infected human macrophages reveals distinct reprogramming signatures. mBio. 2016;7:e00027–16.

31. Westermann AJ, Förstner KU, Amman F, Barquist L, Chao Y, Schulte LN, et al. Dual RNA-seq unveils noncoding RNA functions in host-pathogen interactions. Nature. 2016;529:496–501.

32. Rosani U, Varotto L, Domeneghetti S, Arcangeli G, Pallavicini A, Venier P. Dual analysis of host and pathogen transcriptomes in ostreid herpesvirus 1-positive Crassostrea gigas. Environ. Microbiol. 2015;17:4200–12.

33. Ovington KS, Alleva LM, Kerr EA. Cytokines and immunological control of Eimeria spp. Int. J. Parasitol. 1995;25:1331–51.

34. Stephenson LS, Latham MC, Ottesen EA. Malnutrition and parasitic helminth infections. Parasitology. 2000;121:S23–38.

35. Aloisio F, Filippini G, Antenucci P, Lepri E, Pezzotti G, Cacciò SM, et al. Severe weight loss in lambs infected with Giardia duodenalis assemblage B. Vet. Parasitol. 2006;142:154–8.

36. Preston-Mafham RA, Sykes AH. Changes in body weight and intestinal absorption during infections with Eimeria acervulina in the chicken. Parasitology. 1970;61:417.

37. Sharman PA, Smith NC, Wallach MG, Katrib M. Chasing the golden egg: Vaccination against poultry coccidiosis. Parasite Immunol. 2010;32:590–98

38. Stange J. Studies on host-pathogen interactions at mucosal barrier surfaces using the murine intestinal parasite Eimeria falciformis - Deutsche Digitale Bibliothek. 2012. http://www.deutsche-digitale-bibliothek.de/item/DDAKP5LJSJBAPPALDVQ5Y52YV3AG7NCL. Accessed 16 Dec 2016

39. Kuhn KA, Manieri NA, Liu T-C, Stappenbeck TS. IL-6 stimulates intestinal epithelial proliferation and repair after injury. PloS One. 2014;9:e114195.

40. Park H, Li Z, Yang XO, Chang SH, Nurieva R, Wang Y-H, et al. A distinct lineage of CD4 T-cells regulates tissue inflammation by producing interleukin 17. Nat. Immunol. 2005;6:1133–41.

41. Beck PL, Rosenberg IM, Xavier RJ, Koh T, Wong JF, Podolsky DK. Transforming growth factor-beta mediates intestinal healing and susceptibility to injury in vitro and in vivo through epithelial cells. Am. J. Pathol. 2003;162:597–608.

42. Suzuki A, Sekiya S, Gunshima E, Fujii S, Taniguchi H. EGF signaling activates proliferation and blocks apoptosis of mouse and human intestinal stem/progenitor cells in long-term monolayer cell culture. Lab. Investig. J. Tech. Methods Pathol. 2010;90:1425–36.

43. Kaiser GC, Polk DB. Tumor necrosis factor alpha regulates proliferation in a mouse intestinal cell line. Gastroenterology. 1997;112:1231–40.

44. VanDussen KL, Carulli AJ, Keeley TM, Patel SR, Puthoff BJ, Magness ST, et al. Notch signaling modulates proliferation and differentiation of intestinal crypt base columnar stem cells. Dev. Camb. Engl. 2012;139:488–97.

45. Macian F. NFAT proteins: key regulators of T-cell development and function. Nat. Rev. Immunol. 2005;5:472–84.

46. Reid AJ, Blake DP, Ansari HR, Billington K, Browne HP, Bryant JM, et al. Genomic analysis of the causative agents of coccidiosis in domestic chickens. Genome Res. 2014;gr.168955.113–.

47. Walker R a, Sharman P a, Miller CM, Lippuner C, Okoniewski M, Eichenberger RM, et al. RNA Seq analysis of the Eimeria tenella gametocyte transcriptome reveals clues about the molecular basis for sexual reproduction and oocyst biogenesis. BMC Genomics. 2015;16:1–20.

48. Hehl AB, Basso WU, Lippuner C, Ramakrishnan C, Okoniewski M, Walker RA, et al. Asexual expansion of Toxoplasma gondii merozoites is distinct from tachyzoites and entails expression of non-overlapping gene families to attach, invade, and replicate within feline enterocytes. BMC Genomics. 2015;16:66.

49. Gajria B, Bahl A, Brestelli J, Dommer J, Fischer S, Gao X, et al. ToxoDB: an integrated Toxoplasma gondii database resource. Nucleic Acids Res. 2007;36:D553–6.

50. Reid AJ, Vermont SJ, Cotton JA, Harris D, Hill-Cawthorne GA, Könen-Waisman S, et al. Comparative genomics of the Apicomplexan parasites Toxoplasma gondii and Neospora caninum: Coccidia differing in host range and transmission strategy. PLoS Pathog. 2012;8:e1002567.

51. Gardner MJ, Hall N, Fung E, White O, Berriman M, Hyman RW, et al. Genome sequence of the human malaria parasite Plasmodium falciparum. Nature. 2002;419:498–511.

52. Mineo JR, Kasper LH. Attachment of Toxoplasma gondii to host cells involves major surface protein, SAG-1 (P-30). Exp. Parasitol. 1994;79:11–20.

53. Grimwood J, Smith JE. Toxoplasma gondii: the role of parasite surface and secreted proteins in host cell invasion. Int. J. Parasitol. 1996;26:169–73.

54. Cowman AF, Crabb BS. Invasion of red blood cells by malaria parasites. Cell. 2006;124:755–66.

55. Carruthers V, Boothroyd JC. Pulling together: an integrated model of Toxoplasma cell invasion. Curr. Opin. Microbiol. 2007;10:83–9.

56. Chow Y-P, Wan K-L, Blake DP, Tomley F, Nathan S. Immunogenic Eimeria tenella glycosylphosphatidylinositol-anchored surface antigens (SAGs) induce inflammatory responses in avian macrophages. PLoS ONE. 2011 http://www.ncbi.nlm.nih.gov/pmc/articles/PMC3182191/ Accessed 29 Dec 2016.

57. Tabarés E, Ferguson D, Clark J, Soon P-E, Wan K-L, Tomley F. Eimeria tenella sporozoites and merozoites differentially express glycosylphosphatidylinositol-anchored variant surface proteins. Mol. Biochem. Parasitol. 2004;135:123–32.

58. Talevich E, Kannan N. Structural and evolutionary adaptation of rhoptry kinases and pseudokinases, a family of coccidian virulence factors. BMC Evol. Biol. 2013;13:117.

59. Oakes RD, Kurian D, Bromley E, Ward C, Lal K, Blake DP, et al. The rhoptry proteome of Eimeria tenella sporozoites. Int. J. Parasitol. 2013;43:181–8.

60. Taylor S, Barragan A, Su C, Fux B, Fentress SJ, Tang K, et al. A secreted serine-threonine kinase determines virulence in the eukaryotic pathogen Toxoplasma gondii. Science. 2006;314:1776–80.

61. Saeij JPJ, Coller S, Boyle JP, Jerome ME, White MW, Boothroyd JC. Toxoplasma co-opts host gene expression by injection of a polymorphic kinase homologue. Nature. 2007;445:324–7.

62. Fleckenstein MC, Reese ML, Könen-Waisman S, Boothroyd JC, Howard JC, Steinfeldt T. A Toxoplasma gondii Pseudokinase Inhibits Host IRG Resistance Proteins. PloS Biol. 2012;10:e1001358.

63. Fox BA, Rommereim LM, Guevara RB, Falla A, Triana MAH, Sun Y, et al. The Toxoplasma gondii rhoptry kinome is essential for chronic infection. mBio. 2016;7:e00193–16.

64. Schmid M, Lehmann MJ, Lucius R, Gupta N. Apicomplexan parasite, Eimeria falciformis, co-opts host tryptophan catabolism for life cycle progression in mouse. J. Biol. Chem. 2012;287:20197–20207.

65. R Development Core Team. R: A language and environment for statistical computing. R Foundation for Statistical Computing, Vienna, Austria. 2008. http://www.R-project.org

66. Kowalik S, Zahner H. Eimeria separata: method for the excystation of sporozoites. Parasitol. Res. 1999;85:496–9.

67. Schmatz DM, Crane MSJ, Murray PK. Purification of Eimeria sporozoites by DE-52 anion exchange chromatography. J. Protozool. 1984;31:181–183.

68. MacManes MD. On the optimal trimming of high-throughput mRNA sequence data. Front. Genet. 2014. http://journal.frontiersin.org/article/10.3389/fgene.2014.00013/abstract. Accessed 29 Dec 2016.

69. Trapnell C, Pachter L, Salzberg SL. TopHat: discovering splice junctions with RNA-Seq. Bioinformatics. 2009;25:1105–11.

70. Langmead B, Salzberg SL. Fast gapped-read alignment with Bowtie 2. Nat. Methods. 2012;9:357–9.

71. Liao Y, Smyth GK, Shi W. featureCounts: an efficient general purpose program for assigning sequence reads to genomic features. Bioinformatics. 2014;30:923–30.

72. Robinson MD, McCarthy DJ, Smyth GK. edgeR: a Bioconductor package for differential expression analysis of digital gene expression data. Bioinformatics. 2010;26:139–140.

73. Benjamini Y, Hochberg Y. Controlling the false discovery rate: A practical and powerful approach to multiple testing. J. R. Stat. Soc. Ser. B Methodol. 1995;57:289–300.

